# Odor Blocking Of Stress

**DOI:** 10.1101/2021.02.24.432809

**Authors:** Eun Jeong Lee, Luis R. Saraiva, Naresh K. Hanchate, Xiaolan Ye, Gregory Asher, Jonathan Ho, Linda B. Buck

**Author notes:** **Corresponding author**: Correspondence and requests for materials should be addressed to L.B.B.

## Abstract

Scents have been employed for millennia to allay fear and stress, but whether they do so is poorly understood. In response to fear and stress, hypothalamic corticotropin releasing hormone neurons (CRHNs) induce increases in blood stress hormones. Here, we find that certain structurally and perceptually dissimilar odorants can block mouse stress hormone responses to three potent stressors: physical restraint, predator odor, and male-male social confrontation. Both odorants activate GABAergic inhibitory neurons presynaptic to CRHNs in the hypothalamic ventromedial nucleus (VMH). Stimulation of those neurons inhibits restraint-induced activation of CRHNs and stress hormone increase, mimicking a blocking odorant. Conversely, their silencing prevents odorant blocking of both responses. Notably, we also observed odor blocking of stressor activation in neurons presynaptic to CRHNs in the bed nucleus of the stria terminalis (BNST). Together, these findings indicate that selected odorants can indeed block stress responses, and that odor blocking can occur via two routes: a direct route in which blocking odor signals directly inhibit CRHNs and an indirect route in which they inhibit stressor activation of neurons presynaptic to CRHNs and prevent them from transmitting stress signals to CRHNs.

---

The olfactory system of mammals has evolved to detect a multitude of volatile chemicals that are perceived as odors as well as social cues that elicit instinctive behaviors or physiological effects essential to survival (*1*). The historical use of odors to quell stress (*2*) suggests that some odorants that alone elicit odor perceptions but not instinctive responses might nonetheless dampen innate responses to danger. One hallmark of fear and stress is an elevation of blood stress hormones by corticotropin releasing hormone neurons (CRHNs) in the paraventricular nucleus of the hypothalamus (*3-5*). Here, we investigated whether common odorants can suppress the stress hormone response of mice to three potent stressors: physical restraint, predator odor, and male-male social confrontation.

We selected four common odorants that are attractive to mice, but have dissimilar structures and perceived odor qualities: 2 phenyl ethanol (2PE) (rose) (*6*), trimethylamine (TMA) (fishy) (*7*), hinokitiol (Hino) (woody), and propionic acid (PPA) (cheese) (*7*) (Fig. 1A). Filter paper containing each odorant increased investigation time compared to vehicle, confirming its attractiveness (Fig. 1B). With the exception of PPA, the odorants do not increase plasma levels of the stress hormone ACTH (adrenocorticotropic hormone) (Fig. 1C).

**Fig. 1.**
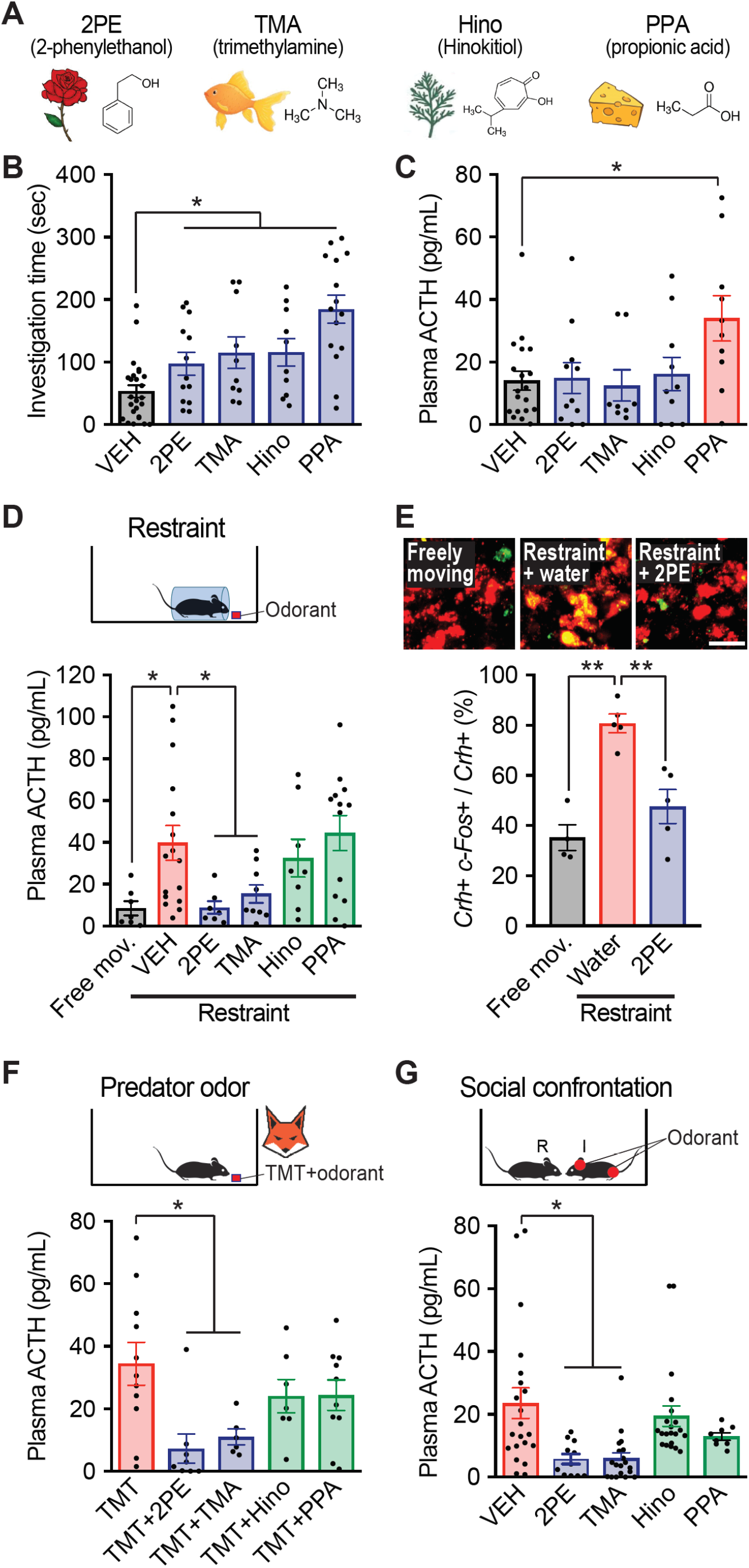
Selected odors block hormonal responses to diverse stressors. Column heights indicate means, error bars indicate S.E.M., and dots in the same column indicate different animals. (A-C) Four odorants with different structures and perceived odors and derivations (A) were attractive to mice compared to vehicle based on investigation time (B). Whereas one (PPA) induced an increase in stress hormone (plasma ACTH), the others did not (C). n = 8–25 per condition. Unpaired *t* test, *P < 0.05. (D-E) Physical restraint caused an increase in stress hormone, which was blocked by exposure to 2PE and TMA, but not Hino or PPA. n = 7–16 per condition. Unpaired *t* test, *P < 0.05 (D). 2PE also blocked restraint-induced activation of CRHNs, as indicated by the percentage of CRHNs labeled for *c-Fos* (E). n = 4–5 per condition. Unpaired *t* test, **P < 0.01. The merged photographic images above show sections costained for CRH (red) and *c-Fos* (green), with colabeled cells indicated in yellow. Scale bar, 20 μm. (F) Predator odor (TMT) exposure stimulated an increase in plasma ACTH that was also blocked by 2PE and TMA, but not Hino or PPA. n = 6–19 per condition. Unpaired *t* test, *P < 0.05, **P < 0.01. G) Social confrontation induced an increase in plasma ACTH when a resident male (R) was exposed to an intruder male (I). Painting the intruder head and genital region with 2PE or TMA, but not Hino or PPA, blocked the stress hormone increase. n = 8–21 per condition. Unpaired *t* test, *P < 0.05.

We found that two of the odorants, 2PE and TMA, block the stress hormone response to physical restraint. Mice were placed in a restrainer that restricts lateral and anterior-posterior movement, but has a hole near the animal’s nose that allows it to smell an odorant placed on a nearby piece of filter paper. Compared to vehicle, 2PE reduced plasma ACTH by 4.5-fold and TMA by 2.6-fold (Fig. 1D) Consistent with this effect, 2PE also decreased the activation of CRHNs by restraint, as indicated by the neural activity marker, *c-Fos* mRNA (Fig. 1E and Fig. S1). Restraint plus 2PE resulted in a 41 ± 11.5 % decrease in CRHNs labeled for *c-Fos* compared to restraint plus water.

We determined that 2PE and TMA also block the stress hormone response to the fox predator odor TMT (2,5-dihydro-2,4,5-trimethylthiazoline) (*8*). Filter paper was placed at one end of the animal’s cage that contained vehicle, TMT, or TMT together with 2PE, TMA, Hino, or PPA. Plasma ACTH seen with TMT alone was decreased 4.8-fold by 2PE and 3.1-fold by TMA (Fig. 1F). These results are reminiscent of a report that rose oil blocks the ACTH response to TMT (*9*), but differ from a reported ability of a higher concentration of Hino to do so (*10*).

Strikingly, 2PE and TMA also inhibit the stress hormone response to male-male social confrontation (*11*). Male mice (“residents”) were singly housed and then exposed to an intruder male whose head and genital region were painted with an odorant or vehicle. Plasma ACTH in residents exposed to intruders painted with vehicle was reduced 4.1-fold when intruders were painted with 2PE and 3.9-fold when they were painted with TMA (Fig. 1G).

How might two common odorants block stress hormone responses to three different stressors? One conceivable possibility is that odor blocking can occur via a direct route. For example, blocking odorants might activate inhibitory neurons presynaptic to CRHNs, which in turn suppress CRHN activation by stressors and their induction of stress hormone increase (Fig. 2A).

**Fig. 2.**
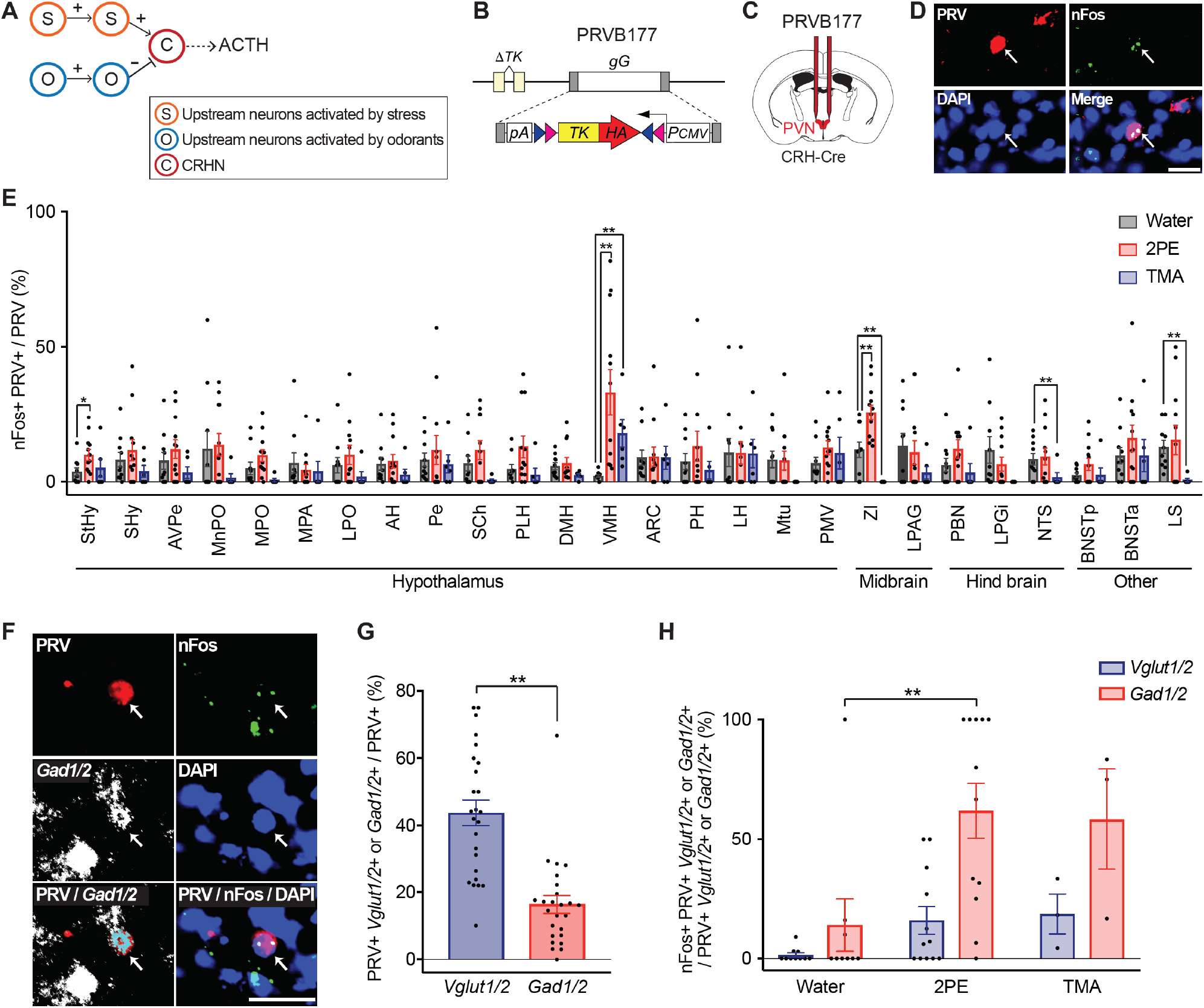
Blocking odors activate inhibitory neurons presynaptic to CRHNs in VMH. (A) In a direct model of odor blocking, blocking odorants activate inhibitory neurons presynaptic to CRHNs, thereby suppressing CRHNs and their activation by stressors. (B-D) PRVB177, which has Cre-dependent expression of HA-TK (B), was used to infect Cre-expressing CRHNs in CRH-ires-Cre mice (C). On d3pi, animals were exposed to water, 2PE, or TMA and brain sections were costained with anti-HA antibodies (PRV) and Fos riboprobes to identify activated (nFos+) PRV+ neurons presynaptic to CRHNs. Nuclei were counterstained with DAPI. Image shows a PRV+ neuron with two nuclear dots of nFos mRNA (arrows) after exposure to 2PE. Scale bar, 20 μm (D). (E) Graph shows percentages of PRV+ neurons with nFos in different brain areas after exposure to water, 2PE, or TMA. Column heights indicate means, error bars indicate S.E.M., and dots in the same column indicate different animals. n = 6–12 per condition. Unpaired *t* test, *P < 0.05, **P < 0.01. PRV+ neurons in the VMH showed significant activation by both 2PE and TMA. (F-H) Following exposure to water, 2PE, or TMA, VMH sections were costained for HA (PRV), nFos, and markers of glutamatergic neurons (*Vglut1/2*) or GABAergic neurons (*Gad1/2*), and then counterstained with DAPI. A photographic image shows one PRV+ neuron costained for nFos and *Gad1/2* (arrows). Scale bars, 20 μm (F). Some VMH PRV+ neurons were colabeled for either *Vglut1/2* or *Gad1/2* (G). Both 2PE and TMA activated *Gad1/2*+ PRV+ neurons in the VMH (H). Column heights indicate means, error bars indicate S.E.M., and dots in the same column indicate different animals. n = 3–12 per condition. Unpaired *t* test, **P < 0.01.

To examine this possibility, we first asked whether 2PE or TMA activates neurons presynaptic to CRHNS. Using CRH-ires-Cre mice (*12*), we infected Cre-expressing CRHNs with PRV177, a Cre-dependent Pseudorabies virus that travels retrogradely across synapses (Fig. 2, B and C) (*3*). On day 3 post-infection (d3pi), when the virus has crossed one synapse, animals were exposed to vehicle, 2PE, or TMA. Brain sections were then costained for PRV (anti-HA antibodies) and the neural activity marker, nuclear Fos mRNA (nFos) (Fig. 2D). As previously, PRV+ neurons presynaptic to CRHNs were seen in numerous brain areas (*3*). However, significant activation of PRV+ neurons by both 2PE and TMA was seen in only one area, the ventromedial nucleus of the hypothalamus (VMH) (Fig. 2E).

Dual staining for PRV and markers of excitatory glutamatergic neurons (*Vglut1* and *Vglut2*) or inhibitory GABAergic neurons (*Gad1* and *Gad2*) indicated that VMH neurons presynaptic to CRHNs can be glutamatergic or GABAergic (Fig. 2, F and G). Triple staining for PRV, nFos, and *Vglut1/2* or *Gad1/2* further showed that both 2PE and TMA can activate VMH GABAergic neurons presynaptic to CRHNs (Fig. 2, F and H, and Fig. S2).

These findings suggested that VMH GABAergic neurons could directly block stress hormone responses by conveying 2PE and TMA odor blocking signals to CRHNs. To examine this possibility, we used chemogenetics to determine whether activation or silencing of VMH GABAergic neurons can alter the stress hormone response and CRHN activation by physical restraint or its blocking by 2PE. Using Gad2-ires-Cre mice (*13*), we infected Gad2+ neurons in the VMH with an AAV that has Cre-dependent expression of the hM3Dq or hM4Di receptor fused to mCherry (Fig. 3, A and E). Upon binding to the synthetic ligand clozapine-n-oxide (CNO), hM3Dq activates neurons whereas hM4Di silences neurons (*14*). Animals were subsequently analyzed for activated CRHNs and increase in stress hormone, and brain sections immunostained for mCherry to determine the locations of neurons expressing hM3Dq or hM4Di (Fig. S3).

**Fig. 3.**
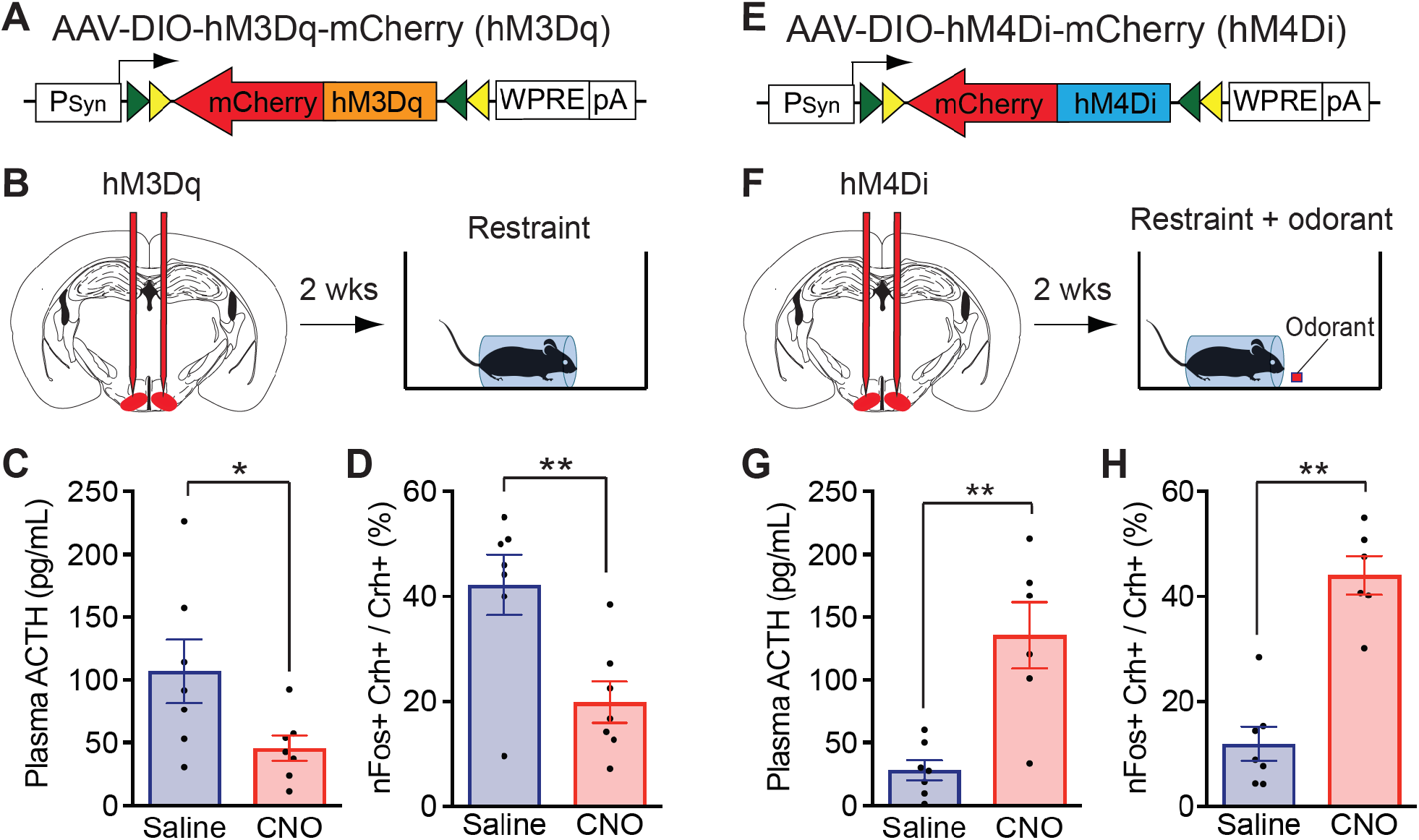
Silencing VMH GABAergic neurons affects odor blocking of stress. (A-B and E-F) AAVs with Cre-dependent expression of hM3Dq or hM4Di were used to activate (hM3Dq) (A) or silence (hM4Di) (E) VMH GABAergic neurons in Gad2-Cre mice. After ∼2 wks, animals were injected with saline, or with CNO to activate hM3Dq or hM4Di, and then exposed to physical restraint (hM3Dq) (B) or physical restraint plus 2PE (F). Plasma ACTH was then measured and brain sections costained for *CRH* and nFos. (C-D) Activation of VMH Gad2+ neurons with hM3Dq inhibits restraint-induced increases in ACTH and CRHN activation. Following restraint, CNO treated animals showed significant decreases in plasma ACTH (C) and in the percentage of CRHNs stained for nFos (D) compared to animals treated with saline. Column heights indicate means, error bars indicate S.E.M., and dots in the same column indicate different animals. n = 7 per condition. Unpaired *t* test, *P < 0.05, **P < 0.01. (G-H) Silencing of VMH Gad2+ neurons with hM4Di interferes with the ability of 2PE to block restraint-induced increases in plasma ACTH and CRHN activation. In animals exposed to restraint plus 2PE, CNO treatment resulted in significant increases in plasma ACTH (G) and the percentage of CRHNs stained for nFos (H) compared to saline treatment. Column heights indicate means, error bars indicate S.E.M., and dots in the same column indicate different animals. n = 6–7 per condition. Unpaired *t* test, **P < 0.01.

Activation of GABAergic VMH neurons by hM3Dq decreased restraint-induced CRHN activation and stress hormone increase. Animals infected with AAV encoding hM3Dq were injected with CNO or control saline, subjected to physical restraint, and plasma ACTH then measured (Fig. 3B). Following restraint, CNO-treated animals showed a 2.4-fold decrease in plasma ACTH relative to saline-treated animals (Fig. 3C). They also showed a 2.1-fold decrease in the percentage of CRHNs activated by restraint, as indicated by nFos mRNA expression (Fig. 3D). These effects of activating VMH GABAergic neurons are similar to those seen with blocking odorant.

Conversely, silencing of VMH GABAergic neurons interfered with the ability of 2PE to block restraint-induced CRHN activation and stress hormone increase. After VMH injection of AAV encoding hM4Di, animals were injected with CNO or saline, exposed to physical restraint in the presence of 2PE, and plasma ACTH then measured (Fig. 3F). Following exposure to restraint and 2PE, CNO-treated animals showed a 4.8-fold increase in ACTH compared to saline-treated controls (Fig. 3G). This was also seen in a few animals expressing hM4Di in an adjacent brain area, suggesting that other areas can also be involved in odor blocking. CNO-treated animals also showed a 3.7-fold increase in the percentage of CRHNs labeled for nFos (Fig. 3H).

Together, these results suggest that odor blocking of stress can occur via a <u>direct route</u> in which blocking odorant activates GABAergic VMH neurons presynaptic to CRHNs, which in turn, suppress CRHNs to prevent their activation by stressors and their induction of a stress hormone response. Although the observed results cannot exclude indirect routes for these effects, the ability of blocking odorants to activate VMH GABAergic neurons presynaptic to CRHNs is highly consistent with a direct route.

Odor blocking of stress could also conceivably occur via an indirect route. In this scheme, blocking odorants would inhibit stressor activation of neurons upstream of CRHNs, thereby preventing those neurons from transmitting stress signals to CRHNs (Fig. 4A). To examine this possibility, we infected CRHNs with PRVB177 and on d3pi exposed animals to restraint or TMT in the presence or absence of 2PE. Brain sections were then costained for HA (PRV) and nFos (Fig. 2D). We previously found that physical restraint and TMT can both activate neurons presynaptic to CRHNs in the anterior bed nucleus of the stria terminalis (BNSTa) (*4*).

**Fig. 4.**
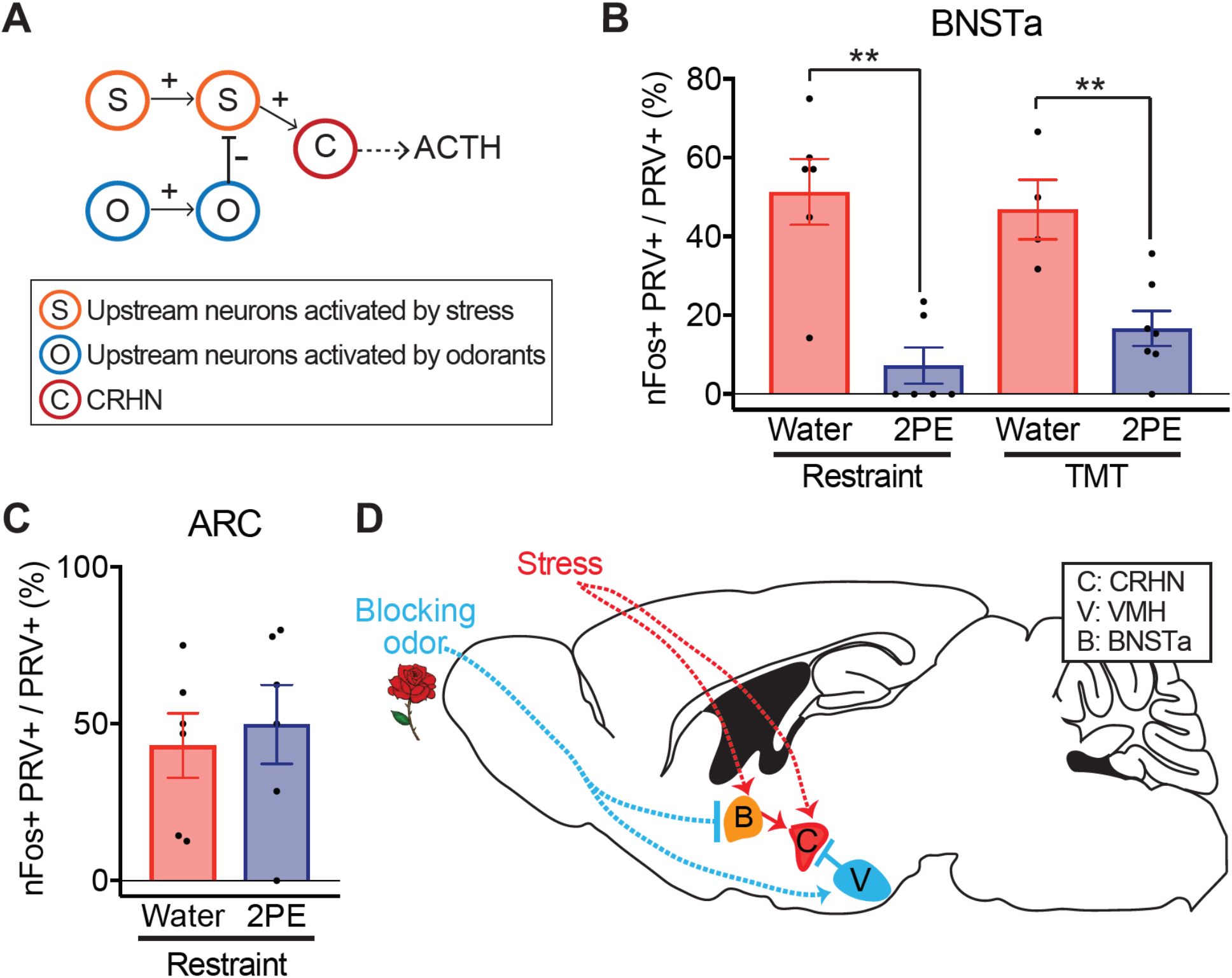
Odor blocking can be indirect. (A) In an indirect model of odor blocking, blocking odors inhibit stressor activation of neurons upstream of CRHNs and prevent them from transmitting stress signals to CRHNs. (B-C) On d3pi of CRHNs with PRVB177, animals were exposed to restraint or TMT in the presence of 2PE or water. Brain sections were then costained for HA (PRV) and nFos mRNA. 2PE significantly decreased the percentage of nFos labeled PRV+ neurons in BNSTa (B). However, 2PE did not significantly affect the percentage of nFos labeled PRV+ neurons in ARC (C). Column heights indicate means, error bars indicate S.E.M., and dots in the same column indicate different animals. n = 4-7 per condition. Unpaired *t* test, **P < 0.01. (D) Schematic showing proposed routes of direct and indirect odor blocking of stress hormone responses to stressors. Signals from blocking odors travel from the nose through the olfactory bulb to the olfactory cortex. From there, they can travel to the VMH, where they activate inhibitory neurons presynaptic to CRHNs and thereby block stressor activation of CRHNs and their induction of ACTH increases. Blocking odor signals can also travel to the BNSTa, where they inhibit stressor activation of neurons upstream of CRHNs and reduce their ability to stimulate CRHNs and ACTH increases.

We found that 2PE inhibits the activation of BNSTa neurons presynaptic to CRHNs by both restraint and TMT (Fig. 4B). Whereas 51.4 ± 8.4% of PRV+ BNSTa neurons were nFos+ following restraint alone, 7.3 ± 4.6% were nFos+ following exposure to restraint plus 2PE. And while 47.0 ± 7.6% of PRV+ BNSTa neurons were nFos+ after exposure to TMT, 16<u>.7</u> ±<u>4.5%</u> were nFos+ after exposure to TMT plus 2PE. In contrast, 2PE had no discernible effect on restraint activation of PRV+ neurons in the ARC (arcuate nucleus of the hypothalamus), which also contains neurons presynaptic to CRHNs (Fig. 4C).

These findings suggest that odor blocking of stress can occur via an <u>indirect route</u> in which blocking odorant inhibits the activation of neurons directly upstream of CRHNs. This inhibition would prevent the upstream neurons from transmitting stressor signals to CRHNs and thereby prevent stressor activation of CRHNs and their ability to induce a stress hormone response.

In these studies, we examined whether odorants can block stress responses and, if so, how they do so. Together, our findings demonstrate that odorants can indeed block physiological stress responses, but only certain odorants have this ability. Our experiments also reveal that odor blocking of stress can occur via direct or indirect pathways that impinge on CRHNs. We identify a direct route in which blocking odorants activate VMH GABAergic inhibitory neurons upstream of CRHNs to inhibit stressor activation of CRHNs and increased stress hormone. We also identify an indirect route in which a blocking odorant inhibits the activation of neurons presynaptic to CRHNs in the BNSTa, thereby preventing the BNSTa neurons from transmitting stressor signals to CRHNs (Fig. 4D).

## ACKNOWLEDGMENTS

We thank members of the Buck laboratory for helpful discussions and comments.

## Author contributions

E.J.L. and L.B.B. conceived the project, E.J.L. conducted experiments and analyzed data, L.S. made initial findings of TMA blocking the ACTH response to TMT, N.K.H and X.Y. analyzed data, G.A. and J.H. performed experiments, and E.J.L and L.B.B. wrote the manuscript.

## Data and materials availability

All data needed to evaluate the conclusions in the paper are present in the paper. Additional data and materials related to this paper will be provided upon reasonable request.

## Materials and Methods

### Mice

Mice aged 2–3 months were used. CRH-IRES-Cre (“CRH-Cre”) mice were generated previously (*12*). C57BL/6J wildtype mice and GAD2-Cre mice (JAX 01082) were purchased from the Jackson Laboratory. All procedures involving mice were approved by the Fred Hutchinson Cancer Research Center Institutional Animal Care and Use Committee. Male mice were used in all experiments. No statistical methods were used to predetermine sample size. Animals were randomly chosen for experimental subjects. Animals were excluded from certain experiments. For nFos experiments shown in Fig. 4, animals were excluded if PRV+ cells were not more than 10 cells in BNSTa. For chemogenetic activation or silencing of the VMH, the animals used had mCherry+ cells indicative of viral infection only in the VMH.

### Viral vectors

#### PRV

PRVB177 was propagated following methods described previously (*3*). Briefly, to propagate PRVB177, PK15 cells (ATCC) were infected with the virus using a multiplicity of infection (m.o.i.) = 0.1–0.01. After infection, cells showed a prominent cytopathic effect (∼2 days). They were harvested by scraping, and the cell material was frozen using liquid nitrogen and then quickly thawed in a 37°C water bath. After three freeze–thaw cycles, cell debris was removed by centrifugation twice at 1,000g for 5 min and the supernatant was then used for experiments. The titre of viral stocks was determined using standard plaque assays on PK15 cells (*15*), with titres expressed in plaque-forming units (p.f.u.).

#### AAVs

Serotype 8 AAVs were used with Cre recombinase-dependent flexstop cassettes that permit expression of mCherry-fused hM3Dq or mCherry-fused hM4Di under the control of the human synapsin promoter (AAV-DIO-hM3Dq-mCherry, AAV-DIO-hM4Di-mCherry) (*16*). The viruses were purchased from the Vector Core at the University of North Carolina at Chapel Hill (the UNC vector core). Amount used is described in virus particles (v.p.).

### Stressors

#### Restraint

Mice were placed individually in a restrainer (a transparent plastic cylinder) (*17*) located in their home cage. The restrainer had a hole in the end near the tip of the animal’s nose. A piece of filter paper (1.5 × 2 cm) with absorbed 50 µL of odorant or VEH (water or DMSO) was placed in the cage near the hole in the cylinder.

#### Predator odor

Mice were exposed to a predator odor or distilled water as described previously (*7*). A piece of filter paper (impregnated with 50 µL of 85 mM 2,5-dihydro-2,4,5-trimethylthiazoline (TMT) (Contech) diluted in water) was dropped gently into one end of the home cage.

#### Social confrontation

Resident male mice were isolated for 10 days before testing, and a male intruder (odorant-swabbed, see below) was placed in a resident’s home cage for 10 min.

### Odor exposure

Olfactory investigation time tests (“olfactory preference test”) were conducted as described previously (*7*), with minor modifications (see below). Adult male C57BL/6J mice were each assayed only once to avoid possible bias attributable to learning. Animals were habituated to the institutional animal facility for at least 5 days after arrival and maintained in single-housed conditions at least 10 days on a 12:12 h light:dark schedule (lights on at 07:00 AM), with experiments performed between 9:00 AM and 11:00 AM. On the day of testing, mice were brought to the experimental room and habituated for 1 hour. In the single odorant experiments, exposure to olfactory stimuli was done by gently dropping a piece of filter paper (1.5 × 2 cm) impregnated with 50 µL of vehicle (VEH, distilled water (H2O) or dimethyl sulfoxide (DMSO)) or odorant (85 mM in VEH, equivalent to 4.25 µmol), into one end of the cage. For the binary odorant mixture exposures, 25 µL of each odorant (4.25 µmol) was placed on a different end of the filter paper for a final concentration of 85 mM of each odorant.

Behavioral tests were video recorded for 15 min with a Canon PowerShot ELPH300HS camera. The total olfactory investigation times were measured during 15 min of odorant exposure.

For ACTH assay in resident male mice confronted with odorant-swabbed intruders, resident male mice were maintained in single-housed conditions at least 10 days on a 12:12 h light:dark schedule (lights on at 07:00 AM), with experiments performed between 5:00 PM and 7:00 PM. On the day of testing, mice were brought to the experimental room and habituated for 1 hour. Exposure to intruders with odorants was done by swabbing with vehicle (VEH, distilled water (H2O) or dimethyl sulfoxide (DMSO)) or odorant (85 mM in VEH, equivalent to 4.25 µmol) on their heads (40 µL) and genitals (10 µL).

For analysis of c-Fos expression in CRH neurons, adult male C57BL/6J mice were exposed to water or 2PE with restraint for 10 min and then perfused 50 min later.

For detection of nFos in PRV-infected cells, CRH-Cre mice injected with PRVB177 3 days earlier were exposed to an odorant (85 mM) with/without stressors for 5 min.

### Stereotaxic injection

Viruses were injected into the brain using a Stereotaxic Alignment System (David Kopf Instruments) with an inhalation anaesthesia of 2.5% isoflurane. Virus suspensions (PRVs: 1–1.5 × 10^6^ p.f.u. (1 µl); AAVs: 1–3 × 10^9^ v.p. (150 nl)) were loaded into a 1-µl syringe, and injected at 100 nl per minute. The needle was inserted into the target location based on a stereotaxic atlas (AP: -0.4, ML: ±0.3, DV: -5.0 mm for PVN; AP: -1.5, ML: ±0.78, DV: -5.75 mm for VMH). After recovery, animals were singly housed with regular 12 h dark/light cycles, and food and water were provided ad libitum.

### Plasma ACTH assay

Plasma ACTH assays were performed as described previously (*3*). Briefly, after mice were killed by cervical dislocation and decapitation, trunk blood was collected directly into blood collection tubes (Becton Dickinson) containing 50 µl aprotinin (Phoenix Pharmaceuticals). Plasma was obtained by centrifugation at 1,600g for 15 min at 4°C, and stored at −80°C. Plasma ACTH concentrations were measured using the ACTH ELISA kit (MD Biosciences), according to the manufacturer’s instructions, with the following modifications: (1) 100 µl of the controls or blood plasma combined with 100 µl of PBS (Phosphate Buffered Saline, pH 7.4) was used in the place of 200 µl plasma and (2) the results were assessed with the QuantaRed Enhanced Chemifluorescent HRP Substrate (Thermo Fisher). Fluorescence was measured with a CytoFluor4000 plate reader (Applied Biosystems).

### In situ hybridization and immunofluorescence

Conventional in situ hybridization was performed essentially as described previously, with some experiments using additional steps for triple staining. Coding region fragments of *Vglut1, Vglut2, Gad1, Gad2, and c-Fos*, and the first intron sequence of *c-Fos* mRNA (for nuclear *c-Fos* (nFos) staining) were isolated from mouse brain cDNA or mouse genomic DNA using PCR, and cloned into the pCR4 Topo vector (Thermo Fisher). Digoxigenin (DIG)- or dinitrophenol (DNP)-labelled cRNA probes (riboprobes) were prepared using the DIG RNA Labelling Mix (Roche) or DNP RNA labeling mix containing DNP-11-UTP (NEL555001EA, Perkin Elmer) and NTPs (Roche). Adult male C57BL/6J mice were perfused transcardially with 4% paraformaldehyde (PFA). Their brains were then soaked in 4% PFA for 4 h, in 30% sucrose for 48 h, and then frozen in OCT (Sakura) and cut into 20 µm coronal sections using a cryostat. Brains of CRH-Cre mice infected 3 days earlier with PRVB177, or wildtype animals, were fresh frozen in OCT, and cut into 20-µm coronal sections using a cryostat. Brain sections were hybridized to DIG- and/or DNP-labelled cRNA probes at 56°C for 13–16 h.

#### Costaining for HA (PRVB177) and nFos mRNA

On d3pi of CRHNs with PRVB177, brain sections were hybridized with riboprobes for *c-Fos* and nFos as described previously (*4*). After hybridization, sections were washed twice in 0.2 × SSC at 63°C for 30 min, incubated with POD-conjugated anti-DIG antibodies (Roche, #11207733910, 1:2,000) and biotinylated anti-HA antibodies (BioLegend, #901505, 1:300) at 37°C for 2 h. Sections were then washed three times for 5 min at RT in TNT buffer, and then treated using the TSA-plus FLU kit (Perkin Elmer). Sections were then washed three times for 5 min at RT in TNT buffer and incubated with 0.5 µg ml^−1^ DAPI and Alexa555-Streptavidin (Thermo Fisher, #32355, 1:1,000) at room temperature for 1 h, and washed. Sections were coverslipped with Fluoromount-G (Southern Biotech).

#### Costaining for HA (PRVB177), nFos mRNA, and Vglut1/2 or Gad1/2 mRNA

On d3pi of CRHNs with PRVB177, sections were hybridized with DIG-labeled riboprobes for nFos and *c-Fos* and a DNP-labeled *Vglut1/2 or Gad1/2* riboprobe. After hybridization, sections were washed twice in 0.2× SSC at 63°C for 30 min, incubated with POD-conjugated anti-DIG antibodies (Roche, #11207733910, 1:2,000), rabbit anti-DNP-KLH antibodies (Molecular Probes, #A6430, 1:200), and biotinylated anti-HA antibodies (BioLegend, #901505, 1:300) at 37°C for 2 h. Sections were then washed three times for 5 min at RT in TNT buffer, and treated using the TSA-plus FLU kit (Perkin Elmer). Sections were then washed three times for 5 min at RT in TNT buffer and incubated with 0.5 µg ml^−1^ DAPI, Alexa555 Streptavidin (Thermo Fisher, #32355, 1:1,000), Alexa647 donkey anti-rabbit IgG (Thermo Fisher, #A21447, 1:1,000) at room temperature for 1 h, and washed. Sections were coverslipped with Fluoromount-G.

RNAscope was conducted using the RNAscope® Multiplex Fluorescent Detection Kit v2 (ACDbio) following the manufacturer’s instructions with minor modifications. Briefly, sections were treated with RNAscope® Protease Reagents for 30 min. After two washes in 1X PBS, sections were hybridized to *Crh* and *c-Fos* probes at 40°C overnight. Bound probes were next amplified by RNAscope® Multiplex FL v2 AMP at 40°C. Sections were then incubated for 15 min at 40°C with RNAscope® Multiplex FL v2 HRPs followed by 30 min of incubation with Opal^2122^ dyes. Sections were next incubated with DAPI and then coverslipped with Prolong Gold Antifade Mountant (Life Technologies).

### Chemogenetic activation and silencin

Chemogenetic experiments were performed as described previously (*4*) with some modifications.

#### Activation

AAV-DIO-hM3Dq-mCherry was injected into the VMH of Gad2-Cre mice by stereotaxic injection (see above). At 2 weeks after injection, mice were intraperitoneally injected with clozapine-N-oxide (CNO; Sigma) (5.0 mg kg^−1^ body weight) in 0.4% DMSO in saline or saline (0.4% DMSO dissolved in saline). Thirty minutes later, mice were exposed to restraint (see above). Trunk blood and brain were collected and the blood was used for plasma ACTH assays (see above). Brains were fixed by soaking in 4% PFA in PBS for 4 h, soaked in 30% sucrose for 24 h, frozen in OCT, and cut into 20-µm coronal sections using a cryostat. Brain sections were washed twice with PBS, permeabilized with 0.5% Triton X-100 in PBS for 5 min, washed twice with PBS, blocked with TNB (Perkin Elmer) for 1 h at room temperature, and then incubated with rabbit anti-RFP (to detect mCherry) (Rockland, #600-410-379, 1:500) diluted in TNB at 4°C overnight. Sections were then washed three times with TNT, incubated with Alexa555 donkey anti-rabbit IgG (Thermo Fisher, #A31572, 1:1,000) and 0.5 µg ml^−1^ DAPI for 1 h at room temperature, and washed three times with TNT. Slides were coverslipped with Fluoromount-G (Southern Biotech).

#### Silencing

AAV-DIO-hM4Di-mCherry was injected into the VMH of Gad2-Cre mice by stereotaxic injection (see above). At 2 weeks after injection, mice were intraperitoneally injected with CNO (5.0 mg kg^−1^ body weight) or saline. Thirty minutes later, mice were exposed to restraint combined with odor exposure to 2PE (see above). Trunk blood and brain were then collected. Blood was used for plasma ACTH assays and brains were treated and immunostained with rabbit-anti-RFP antibody (see above).

### Cell counting

Cell counting was performed as described previously (*4*). Images were collected using an AxioCam camera and AxioImager.Z2 microscope with an apotome device (Zeiss). Images were acquired with auto-exposure setting, because background between different sections and animals is different. No additional post-processing was performed on any of the collected images for counting. Counting was conducted blindly. Brain structures were identified microscopically and in digital photos using a mouse brain atlas (*18*). Every fifth section was analyzed for all experiments. For the data shown in Fig. 2 and Fig. 4, cells were counted in a given area in only one hemisphere, the hemisphere that contained the most PRV+ cells. The percentage of PRV+ cells with nFos labeling among all PRV+ cells lacking cytoplasmic *Fos* labeling was calculated.

### Statistical analysis

All data are shown as the mean ± S.E.M.. Data were tested with the Shapiro–Wilk test for normality. For data with a normal distribution, the unpaired t-test was used to compare two groups to analyze statistical significance. All tests were two-sided.

## Abbreviations for brain areas

Abbreviations used for brain areas are according to our previous reports (*3-5*).

AH: anterior hypothalamic area
ARC: arcuate hypothalamic nucleus
AVPe: anteroventral periventricular nucleus
BNSTa: bed nucleus of the stria terminalis, anterior part
BNSTp: bed nucleus of the stria terminalis, posterior part
DMH: dorsomedial hypothalamic nucleus
LH: lateral hypothalamic area
LPAG: lateral periaqueductal grey
LPGi: lateral paragigantocellular nucleus
LPO: lateral preoptic area
LS: lateral septal nucleus
MnPO: median preoptic nucleus
MPA: medial preoptic area
MPO: medial preoptic nucleus
MTu: medial tuberal nucleus
NTS: nucleus of the solitary tract
PBN: parabrachial nucleus
Pe: periventricular nucleus of the hypothalamus
PH: posterior hypothalamic nucleus
PLH: peduncular part of lateral hypothalamus
PMV: premammillary nucleus, ventral part
SCh: suprachiasmatic nucleus
SHy: septohippocampal nucleus
StHy: striohypothalamic nucleus
VMH: ventromedial hypothalamic nucleus
ZI: zona incerta

---

**Fig. S1.**
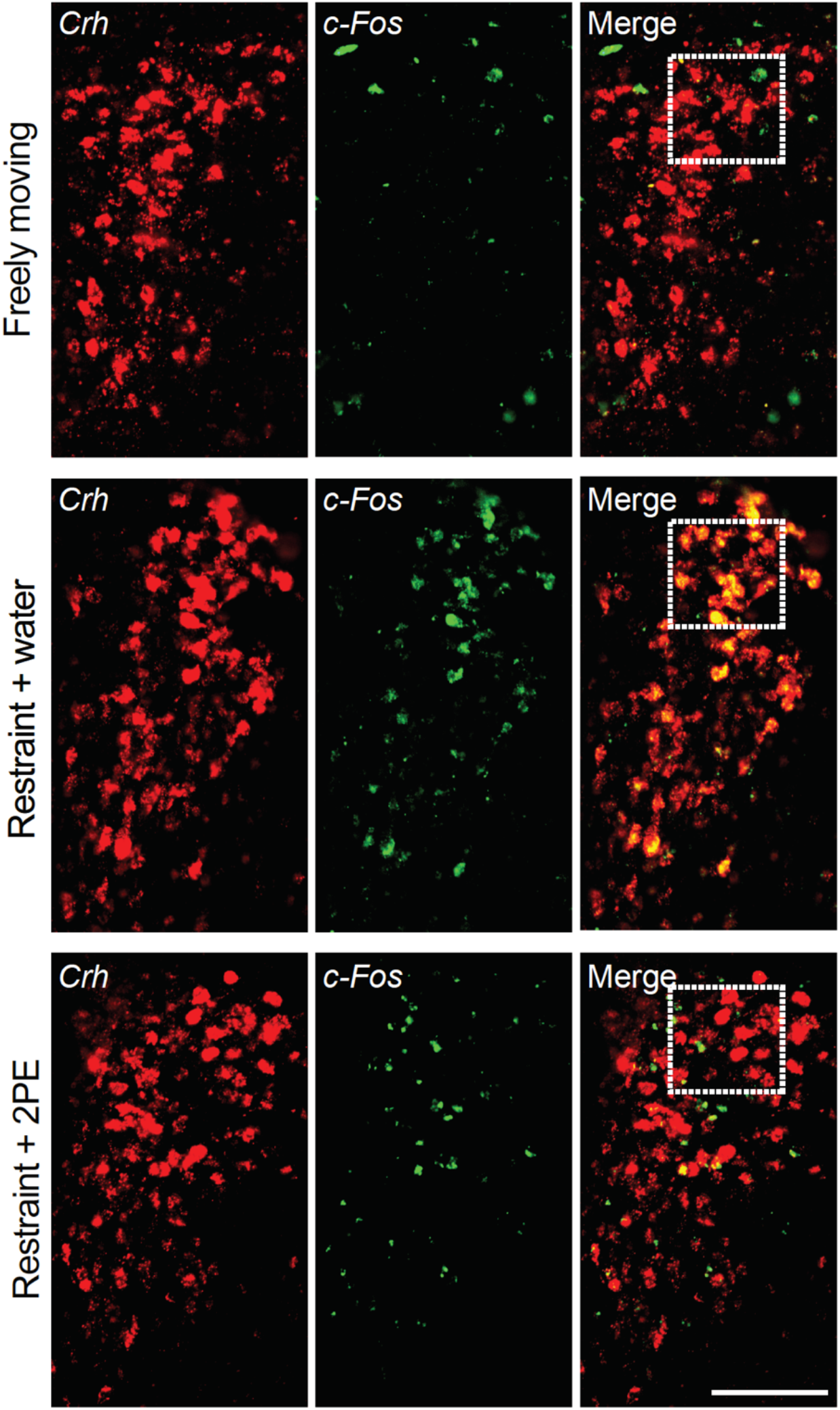
Activation of CRHNs by physical restraint. Brain sections were costained for *Crh* and *c-Fos* mRNAs in freely moving animals, or following exposure to physical restraint plus water or physical restraint plus 2PE. Shown here are photographs of the paraventricular nucleus of the hypothalamus in each case with *Crh*+ neurons shown in red and *c-Fos*+ neurons in green. Merged images are shown at right. Boxes indicate segments shown at higher magnification in Fig. 1E. Scale bar, 100 um.

**Fig. S2.**
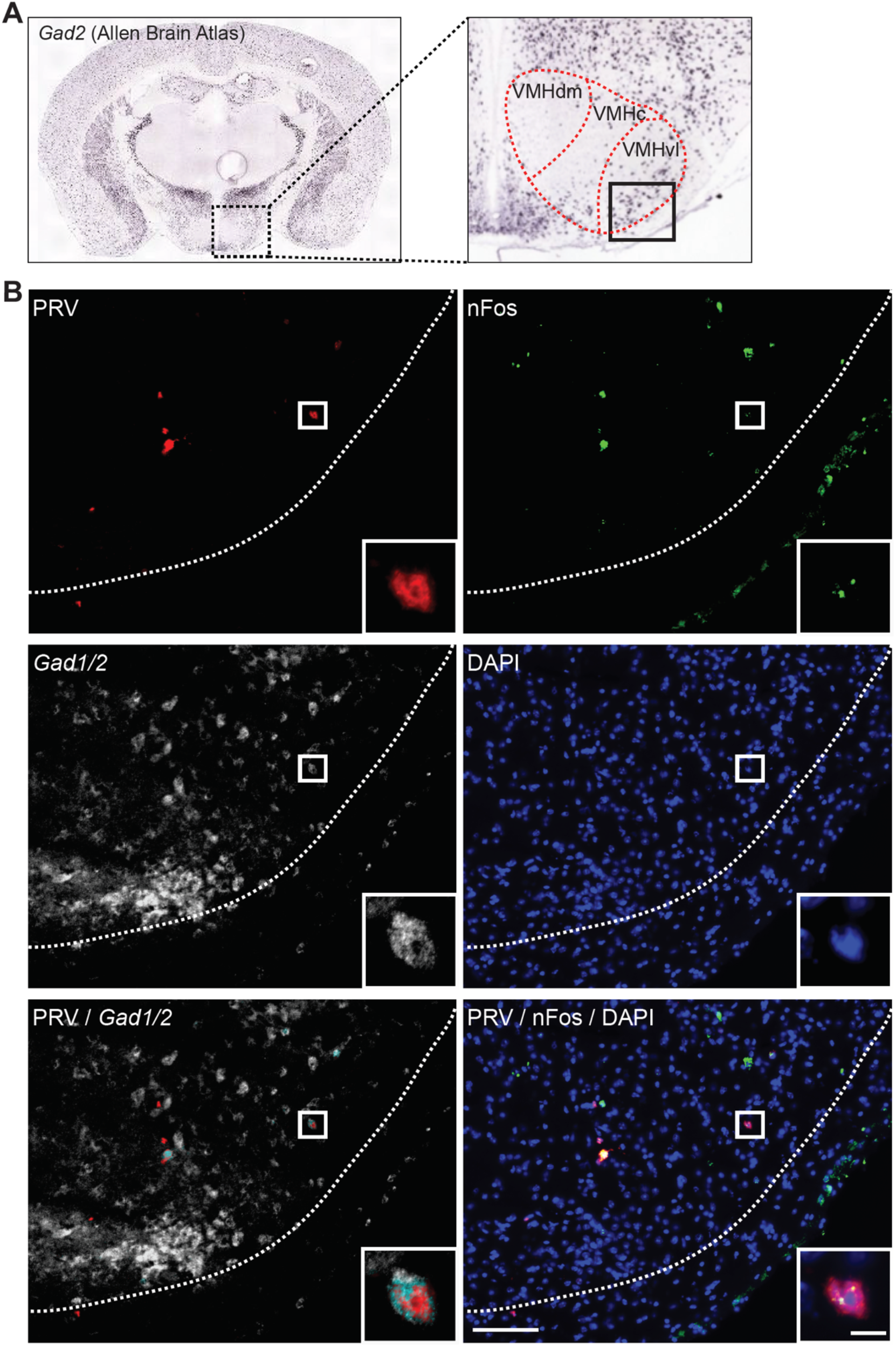
Activation of presynaptic neurons in the VMH by 2PE. A) An image from the Allen Brain Atlas (https://mouse.brain-map.org/) shows staining for *Gad2* mRNA in the VMH. The dotted box at left is shown at higher magnification on the right with the VMH and its subregions outlined in red. The boxed area at right is shown in the photographic images below. B) On day 3 after infection of CRHNs with PRVB177, animals were exposed to 2PE. Brain sections were then costained for PRV (HA) and *Gad1/2* and nFos mRNAs, and counterstained with DAPI. Shown here are images of one VMH section showing labeling for PRV, nFos, Gad1/2, and DAPI. Merged images are also shown for PRV and Gad1/2 and for PRV, nFos, and DAPI. The boxed area is shown at higher magnification in the lower right corner of each image. The cell shown was labeled for PRV, nFos, and Gad1/2. Scale bars, 100 um (left), 10 um (right).

**Fig. S3.**
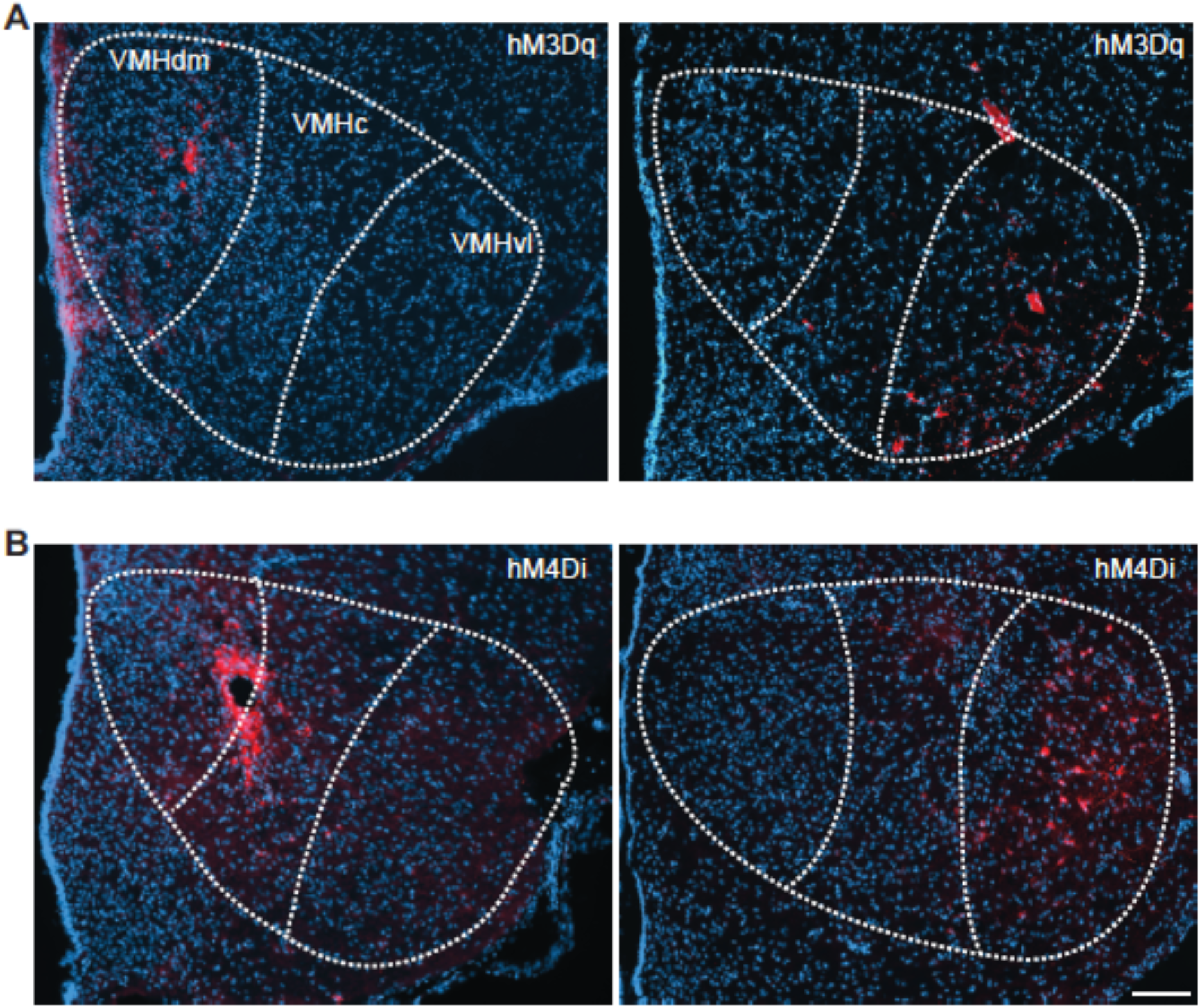
Locations of VMH neurons infected with an activating or silencing virus. The VMH was injected with an AAV with Cre-dependent expression of hM3Dq-mCherry (A) or hM4Di-mCherry (B). Approximately two weeks later, brain sections were stained with antibodies that recognize mCherry (red) and counterstained with DAPI (blue). The photographic images here show mCherry+ cells in the VMH of two animals injected with each virus. The VMH and its subregions are outlined with dotted lines. Scale bar, 100 um.

